# Evaluation design shapes estimates of cross-donor generalization in single-cell classification of cell-type

**DOI:** 10.64898/2026.10.26.761949

**Authors:** Xingyi Chen, Jiabei Li, Alexis Battle

## Abstract

**Motivation:** Single-cell cell-type classifiers are often evaluated using random cell-level splits, although cells from the same donor are not independent. We asked how evaluation design affects estimates of cross-donor generalization.

**Results:** Across PBMC and lung datasets, random cell-level splitting consistently yielded higher macro F1 than donor-held-out evaluation across multiple expression representations. This gap persisted after batch covariate adjustment and donor-level mixed-effects modeling. In contrast, evaluation gaps were generally minimal in highly separable, trivially easy kidney, pancreas, and blood classification settings and in a randomized-label setting with no meaningful predictive signal. These results show that evaluation bias depends on the underlying classification difficulty and that donor-held-out evaluation is necessary when claims concern generalization to unseen individuals.

**Availability and Implementation:** Code and analyses are available on GitHub at https://github.com/danielchen05/scRNA-cross-donor-generalization/. Processed data are archived at Zenodo at https://doi.org/10.5281/zenodo.19717558.

## 1. Introduction

Single-cell RNA sequencing (scRNA-seq) has enabled increasingly detailed characterization of cellular heterogeneity across tissues, disease states, and individuals [1]. As the scale of single-cell studies has grown, automated cell-type annotation has become central to scRNA-seq analysis. Supervised approaches can transfer labels from annotated reference datasets to new cells, reducing the time and subjectivity associated with manual annotation [2,3]. A central requirement for these methods is generalization: models trained on a reference cohort should ideally maintain reliable performance when applied to cells from new individuals not represented during training. However, it remains unclear how the difference between cell-level and donor-level performance estimates varies across tasks of differing difficulty and whether accounting for donor-associated and technical heterogeneity reduces this gap. Here, we address these questions by directly comparing the two evaluation strategies across five cohorts using a common classifier and fixed expression representations.

A key challenge in evaluating cross-donor generalization arises from the hierarchical structure of single-cell data. Cells sampled from the same donor share genetic background, environmental exposures, biological state, and often technical processing [4,5]. The consequences of treating cells as independent biological replicates have been established in single-cell differential expression analysis [6]. In supervised cell-type annotation, several benchmarks have included intra-dataset evaluations that partition individual cells without explicitly grouping them by donor. Examples include five-fold cross-validation across cells [3,7] and repeated training and test splits stratified by cell type [8]. Such partitions do not enforce donor separation, and cells from the same individual can potentially appear in both training and test sets. Performance measured in this setting therefore does not, by itself, establish generalization to individuals excluded from classifier training.

Previous studies have also evaluated transfer across independent datasets or sequencing platforms [3,7,8]. These designs assess generalization under changes in dataset composition or measurement conditions, but do not isolate the effect of replacing cell-level partitions with donor-held-out partitions within the same cohort. Donor-separated evaluation also has direct precedents in cell-type annotation: scTab partitions donors into disjoint training, validation, and test sets [9]. A recent preprint additionally examines donor-aware evaluation for donor-level inflammatory bowel disease classification, a distinct prediction task [10]. However, the extent to which random cell-level splitting yields different performance estimates from donor-held-out evaluation within the same dataset remains unclear, which we investigate in detail here.

An additional open question is how the magnitude of this difference varies across datasets with different classification characteristics. Cell-type classification tasks vary considerably in their difficulty and biological resolution. Broad, transcriptionally distinct populations may be classified with near-perfect accuracy regardless of how the data are partitioned, whereas poorly separable populations may remain difficult under either evaluation strategy. Between these extremes, donor-associated variation may have a greater opportunity to influence apparent classification performance. Consequently, the difference between random cell-level and donor-held-out evaluation may depend on the underlying classification difficulty. Establishing when this difference becomes detectable is important for interpreting benchmarks across datasets rather than assuming that an effect observed in one dataset will have the same magnitude in another.

Another unresolved question is whether the evaluation gap persists after accounting for donor-associated and technical heterogeneity. Integration approaches such as Harmony and scVI aim to construct representations that account for unwanted sources of variation [11,12]. Technical covariates can also be included in classifiers, while mixed-effects models can account for donor-associated dependence when analyzing classification outcomes. These approaches serve different purposes, and none substitute for explicitly separating donors between classifier training and testing. It therefore remains important to determine whether differences between evaluation strategies persist across representations and after adjustment for measured technical and donor-associated variation.

Here, we examine how evaluation design influences estimates of cross-donor cell-type classification performance in scRNA-seq data. We compare random cell-level and donor-held-out evaluation across five cohorts spanning different tissues, cohort sizes, and levels of classification difficulty, using a common classifier and fixed, cohort-level expression representations. We further examine batch-aware representations, explicit technical covariates, and donor-level mixed-effects analyses of classification correctness. Together, these analyses clarify how evaluation design and classification difficulty shape estimates of cross-donor generalization, highlighting the importance of matching evaluation strategies to the intended prediction setting.

## 2. Results

### 2.1 Benchmark design for evaluating cross-donor generalization

First, we establish a benchmarking framework. We compared random cell-level splitting with donor-held-out cross-validation across a common cell-type classification framework (**Figure 1**). Under the cell-level split, cells from the same donors could appear in both training and test sets, while donor-held-out evaluation restricted each donor entirely to one partition (**Figure 1A**).

**Figure 1:**
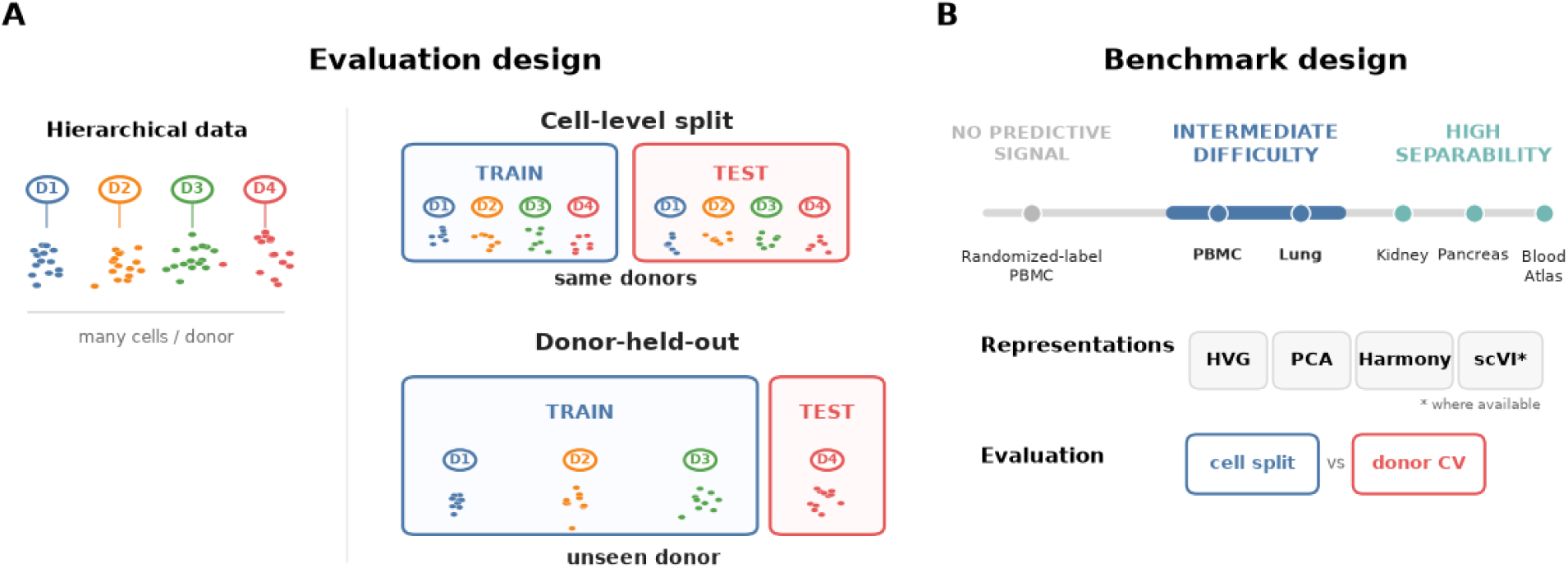
Study design for evaluating cross-donor generalization in single-cell classification. (A): Schematic of the hierarchical structure of scRNA-seq data and the two evaluation strategies compared throughout the study. Random cell-level splitting places cells from the same donors in both training and test sets, whereas donor-held-out cross-validation evaluates performance on cells from individuals excluded from classifier training. (B): Benchmark design spanning classification tasks of varying difficulty. Randomized-label PBMC represents a setting with no meaningful predictive signal; PBMC and lung represent intermediate-difficulty settings; and kidney, pancreas, and Blood Atlas represent highly separable settings. Positions along the continuum are conceptual rather than quantitative. Models were evaluated across HVG, PCA, Harmony, and, where available, scVI representations, comparing random cell-level splitting with donor-held-out cross-validation.

The benchmark included classification tasks spanning a range of difficulty, from highly separable cell types to intermediate settings and a randomized-label setting with no meaningful predictive signal (**Figure 1B**). PBMC and lung datasets represented intermediate settings, while kidney, pancreas, and Blood Atlas datasets represented highly separable classification settings.

Randomizing the PBMC cell-type labels created the setting with no meaningful predictive signal. Across datasets, we evaluated highly variable gene (HVG) expression, PCA, Harmony, and scVI representations where available, comparing each under the two evaluation strategies. This design allowed us to separate the effect of evaluation strategy from representation choice.

### 2.2 Random cell-level splitting overestimates cross-donor performance despite batch adjustment and donor-level modeling

We first evaluated the effect of splitting strategy in the PBMC dataset, which contained 10,107 cells from 23 healthy donors spanning 13 retained cell types (**Figure 2A**). Cell-type abundance and donor representation varied across populations, although all retained cell types satisfied the prespecified support criteria (**Figure S1A**). The resulting dataset contained well-supported transcriptional populations but also detectable structure associated with the collection site, emphasizing that cells from the same donor or study context are not independent observations (**Figure S2A and S3A**).

**Figure 2:**
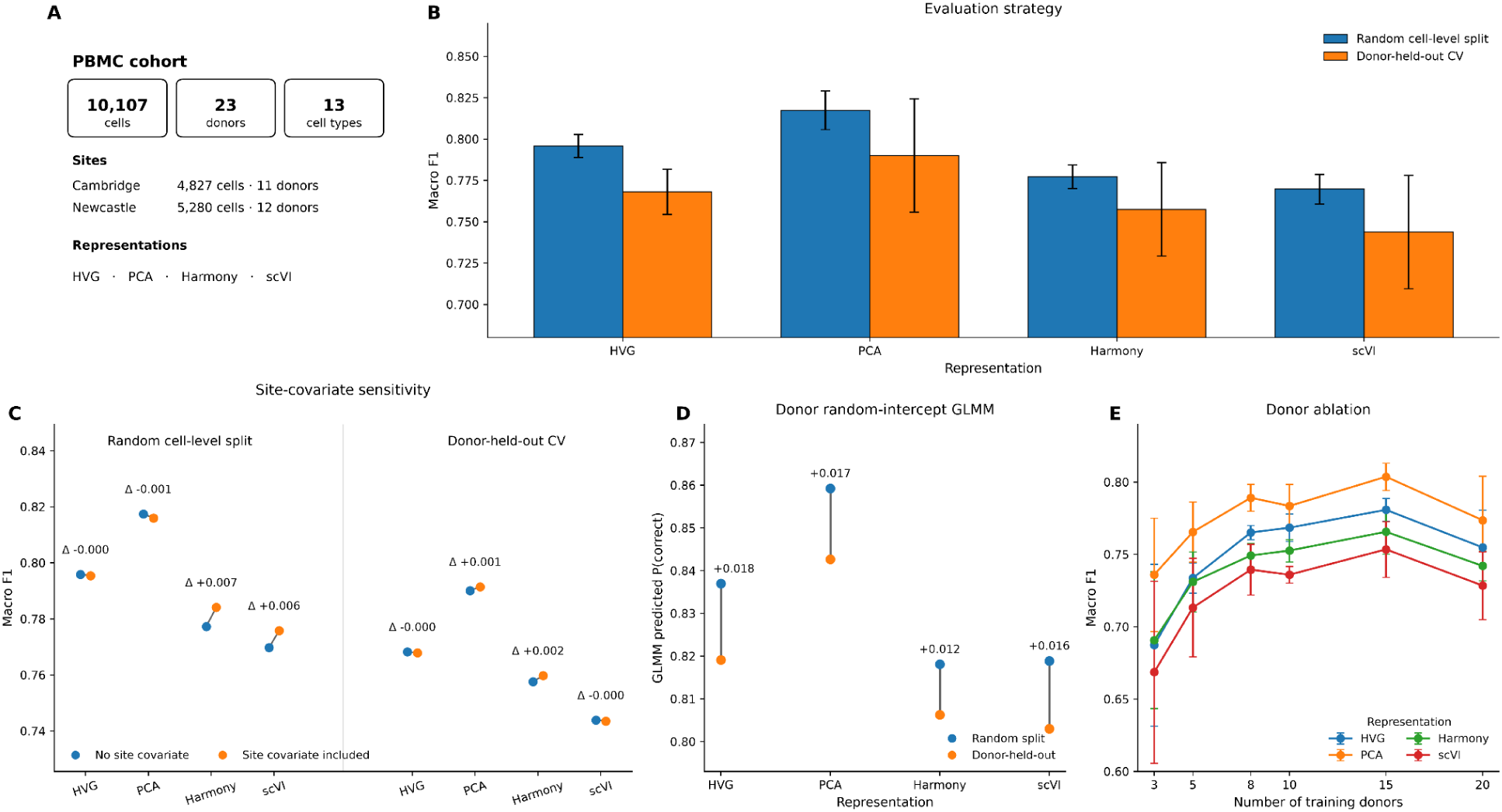
Random cell-level splitting overestimates cross-donor classification performance in the PBMC cohort. (A): Overview of the PBMC cohort, including the number of cells, donors, cell types, study sites, and evaluated expression representations. (B): Macro F1 under random cell-level splitting and donor-held-out cross-validation across HVG, PCA, Harmony, and scVI representations. Bars show mean macro F1, with error bars indicating ± SD across five random cell-level splits or five donor-held-out cross-validation folds, respectively. (C): Sensitivity analysis assessing the effect of including study site as an additional covariate under both evaluation strategies; annotated values show macro F1 with the site covariate minus macro F1 without it, within each evaluation strategy. (D): Donor random-intercept generalized linear mixed model estimates of the probability of correct classification under random cell-level splitting and donor-held-out evaluation, showing that the performance gap persists after accounting for donor-level heterogeneity. (E): Donor-ablation analysis showing macro F1 as a function of the number of training donors, with evaluation on all remaining donors. Points and error bars show the mean ± SD across five donor subsamples at each training-donor count.

Across all four representations, random cell-level splitting produced higher macro F1 than evaluation on held-out donors. The difference was 0.028 for HVG features, 0.027 for PCA, 0.020 for Harmony, and 0.026 for scVI (**Figure 2B**). Thus, the discrepancy was not specific to a particular latent representation and remained after applying methods designed to reduce technical or donor-associated variation. These results indicate that allowing cells from the same donors to appear in both training and test sets yields systematically over-optimistic estimates in comparison to evaluation on entirely unseen donors.

We next examined whether batch effects, specifically the technical effect induced by different data collection sites, could account for this difference. Explicitly adding site as a classifier covariate only produced small changes in performance, with changes in macro F1 of no more than 0.007, and did not eliminate the separation between random cell-level and donor-held-out evaluation (**Figure 2C**). Consistent with this result, modeling cell-level correctness using generalized linear mixed models (GLMMs) while including a donor-level random intercept continued to show higher predicted correctness under random cell-level evaluation across representations (**Figure 2D**). Together, these analyses suggest that the observed discrepancy could not be explained solely by batch effects, specifically the measured collection-site variable.

We also asked whether the gap reflected limited training donor diversity. Increasing the number of donors available for training generally improved performance on unseen donors, demonstrating that cross-donor generalization benefits from broader donor representation (**Figure 2E**). However, the distinction between evaluation strategies remained conceptually separate from the sample-size effect: random cell-level splitting evaluates cells in the presence of donor contexts already represented during training, while donor-held-out evaluation directly measures the transfer to new individuals. As an external-cohort stress test, bidirectional transfer between the Cambridge and Newcastle PBMC cohorts showed lower performance and a persistent directional asymmetry across representations, even after matching cell type support or donor counts (**Figure S4**). These findings reinforce that performance depends on the biological and technical contexts represented during training, and that random cell-level splitting can obscure this dependence.

### 2.3 Overestimation persists in an independent lung atlas

We next asked whether the difference between random cell level and donor-held-out evaluation generalized to a larger, independently assembled dataset. The Lung Atlas benchmark contained 83,966 cells from 104 donors spanning 31 retained cell types and 13 source datasets (**Figure 3A**). Cell populations varied substantially in abundance and donor support, while visualization of retained cells showed both distinct cell-type structure and substantial mixing among related populations (**Figure S1B, S2B**). The data also contained structure associated with the source dataset, providing a more heterogeneous data landscape than the PBMC cohort (**Figure S3B**).

**Figure 3:**
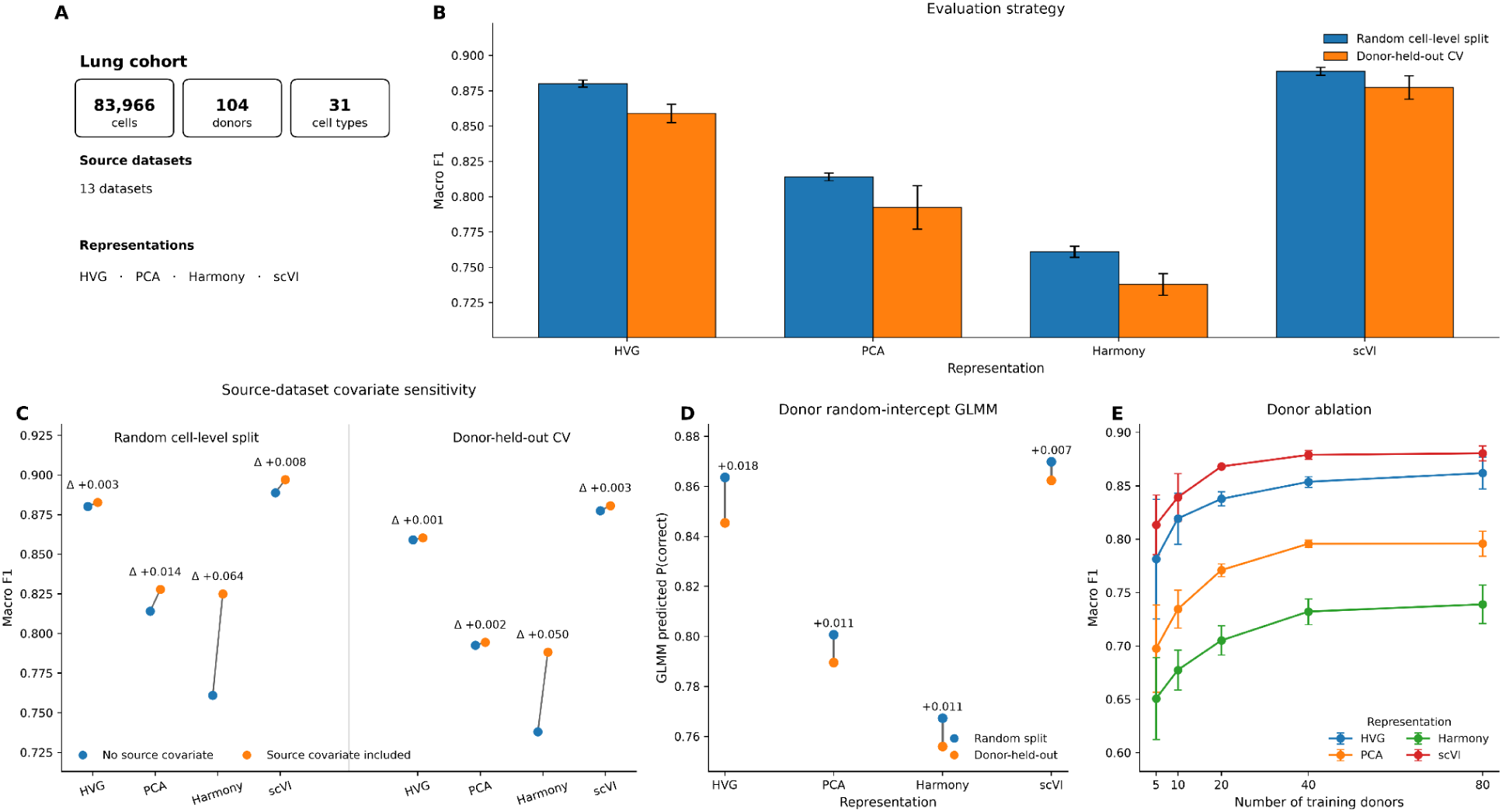
Random cell-level splitting overestimates cross-donor classification performance in the lung cohort. (A): Overview of the lung cohort, including the number of cells, donors, cell types, source datasets, and evaluated expression representations. (B): Macro F1 under random cell-level splitting and donor-held-out cross-validation across HVG, PCA, Harmony, and scVI representations. Bars show mean macro F1, with error bars indicating ± SD across five random cell-level splits or five donor-held-out cross-validation folds, respectively. (C): Sensitivity analysis assessing the effect of including source dataset as an additional covariate under both evaluation strategies; annotated values show macro F1 with the source-dataset covariate minus macro F1 without it, within each evaluation strategy. (D): Donor random-intercept generalized linear mixed-model estimates of the probability of correct classification under random cell-level splitting and donor-held-out evaluation, showing that the performance gap persists after accounting for donor-level heterogeneity. (E): Donor-ablation analysis showing macro F1 as a function of the number of training donors, with evaluation on all remaining donors. Points and error bars show the mean ± SD across five donor subsamples at each training-donor count.

Despite these differences, random cell-level splitting again produced higher macro F1 than donor-held-out evaluation for every representation tested (**Figure 3B**). Macro F1 decreased from 0.880 to 0.859 for HVG features, 0.814 to 0.793 for PCA, 0.761 to 0.738 for Harmony, and 0.889 to 0.877 for scVI when evaluation was changed from random cell level splitting to donor-held-out evaluation. The corresponding differences ranged from 0.011 to 0.023, reproducing the direction and magnitude of the PBMC result in an independent atlas-scale cohort. Notably, the effect was again observed across the board for both conventional dimensionality reduction approaches and representations explicitly designed to account for technical variations.

As the Lung Atlas combines data from multiple studies, we next tested whether source dataset information accounted for the discrepancy. Adding source dataset as a covariate in the classifier altered absolute performance but did not remove the advantage of random cell-level evaluation (**Figure 3C**). The difference in performance between evaluation strategies remained across all four representations after covariate adjustment. Therefore, incorporating source heterogeneity did not explain the evaluation gap.

We obtained the same qualitative conclusion when donor-to-donor heterogeneity was modeled directly in a GLMM. Predicted classification success remained higher under random cell-level evaluation across representations (**Figure 3D**). Thus, accounting statistically for donor-associated variation did not make random cell-level evaluation equivalent to testing on previously unseen individuals.

Finally, we examined how donor-held-out performance changed as the diversity of the training cohort increased. Performance improved rapidly as additional training donors were introduced, and then showed diminishing gains once sufficient numbers of donors were represented (**Figure 3E**). This observation indicates that broader donor sampling improves generalization, but does not eliminate the need to evaluate entirely unseen donors when the goal is to measure cross-donor performance. Together with previous analyses on PBMC, these results reveal that optimism introduced by random cell-level splitting is reproducible across datasets with different scale, tissue context, cell-type complexity, and technical composition.

### 2.4 Evaluation gaps diminish for very easy or very hard classification tasks

Having observed consistent differences between random cell level and donor-held-out evaluation in PBMC and Lung, we next examined settings in which the choice of evaluation strategy was expected to have less influence on the classification performance. Across Kidney, Pancreas, and Blood Atlas datasets, cell type classification was already highly accurate under both evaluation schemes, while randomizing the PBMC cell-type labels provided a complementary setting in which no biologically meaningful predictive signal was present. These additional datasets allowed us to examine evaluation gaps for highly separable cell types and for randomized labels with no meaningful predictive signal (**Figure 4A**).

**Figure 4:**
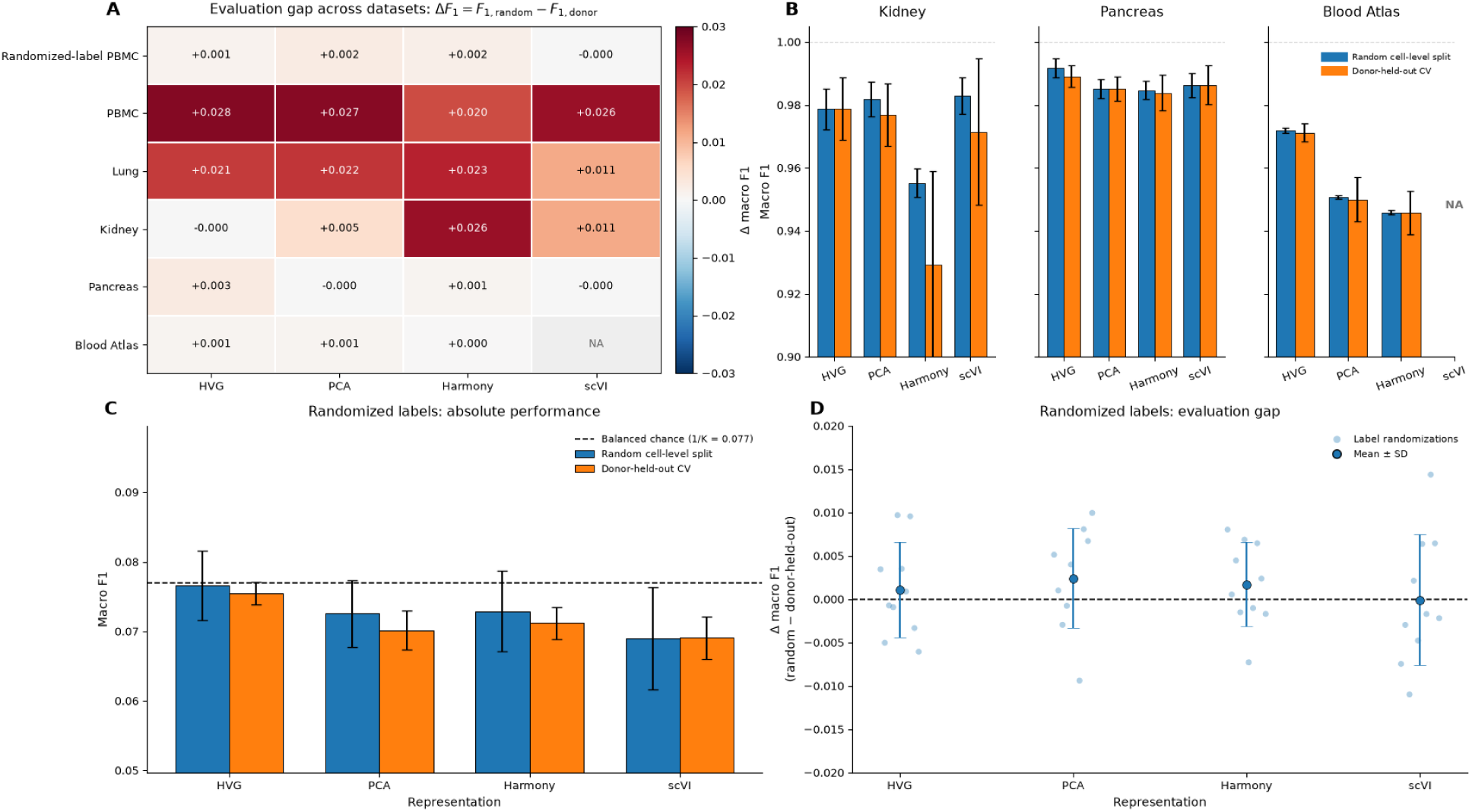
Evaluation gaps across classification tasks of varying difficulty. (A): Heatmap of the evaluation gap, defined as mean macro F1 under random cell-level splitting minus mean macro F1 under donor-held-out cross-validation, across datasets and expression representations. The randomized-label PBMC row shows the mean paired gap across ten label randomizations. Positive values indicate higher performance under random cell-level splitting. NA denotes the unavailable scVI representation for Blood Atlas. (B): Absolute macro F1 in the kidney, pancreas, and Blood Atlas cohorts. Bars and error bars show the mean ± SD across five random cell-level splits or five donor-held-out folds, respectively. Dashed lines mark perfect classification. (C): Absolute macro F1 in the randomized-label PBMC setting, generated by replacing cell-type labels with thirteen donor-balanced random pseudo-classes while retaining the original expression profiles and representations. Bars and error bars show the mean ± SD across ten independent label randomizations. Each randomization was evaluated using one random cell-level split and five-fold donor-held-out cross-validation. The dashed line marks the balanced-chance reference of 1/13, approximately 0.077. (D): Paired random-split minus donor-held-out macro F1 differences for the same label randomizations. Light points show individual randomizations, and darker points with error bars show the mean ± SD. The dashed line marks zero difference.

Kidney and pancreas represented highly separable classification settings in which most representations achieved near-perfect performance. In Pancreas, macro F1 remained 0.98-0.99 under both evaluation regimes, and the corresponding evaluation gaps were effectively zero across representations. Kidney showed similarly high performance for HVG and PCA, with small differences between evaluation schemes, although larger gaps remained for Harmony and scVI (**Figure 4B**). The Blood Atlas provided another highly separable classification setting at atlas scale: despite containing 166 donors, random cell level and donor-held-out performance were almost indistinguishable for HVG, PCA, and Harmony, with macro F1 differences of approximately 0.001 or less (**Figure 4A-B**). The retained cell type composition and donor support of those datasets are shown in **Figure S1C-E**, with transcriptional and technical structure visualized in **Figure S2C-E** and **Figure S3C-E**.

These highly separable settings reveal that random cell-level splitting does not always produce a large apparent advantage. When cell types are already highly separable across donors, both evaluation strategies can yield similarly high performance, leaving little room for difference. The magnitude of the evaluation gap depends on the underlying difficulty of the classification problem rather than being fixed as a consequence of the splitting scheme.

We next examined a randomized-label setting to assess performance when no meaningful predictive signal is present. Cell-type labels in the PBMC dataset were replaced with 13 donor-balanced random pseudo-classes while preserving the original expression profiles and representations. Across 10 independent label randomizations, pseudo-class counts differed by at most one cell within each donor, preventing the impact of donor-specific label imbalance (**Figure S5A**). As expected, macro F1 remained close to the balanced chance-level of 1/13 across representations and evaluation strategies (**Figure 4C, S5B**). Consistent with this near-chance performance, the differences in macro F1 when comparing two strategies were centered close to zero (**Figure 4D**). Therefore, when labels contain no reproducible signal, random cell-level splitting itself does not generate the systematic performance advantage observed in PBMC and Lung.

Together, the highly separable and randomized-label settings illustrate conditions under which the evaluation gap becomes small. The discrepancy is minimal when the classification is either almost perfect across donors or no better than chance, while it is most evident in intermediate settings where biologically meaningful signals coexist with donor-associated variations.

Therefore, these results support interpreting the evaluation gap as an interaction between dataset structure and evaluation design, rather than as a fixed property of single-cell classification tasks.

## 3. Discussion

We show that evaluation design can materially affect estimates of cross-donor cell-type classification performance in scRNA-seq data. In both PBMC and lung, random cell-level splitting consistently produced higher performance than donor-held-out evaluation across multiple representations, including after technical covariate adjustment. Donor-level mixed-effects analyses of classification correctness also supported this difference. By contrast, the gap was small in highly separable tasks with near-perfect performance and in the randomized-label setting with near-chance performance, indicating that its magnitude depends on the underlying classification difficulty rather than being fixed across datasets.

These findings emphasize that accounting for heterogeneity and defining the intended unit of generalization are separate issues. Integration methods, technical covariates, and mixed-effects models may reduce or model structured variation, but they do not make cell-level evaluation equivalent to testing on unseen donors. For studies making claims about performance on new individuals, train-test partitioning should therefore occur at the donor or patient level, and the biological unit used for validation should be reported explicitly. For example, a model intended to annotate immune cells from newly recruited patients should be evaluated on patients excluded from classifier training. Similarly, extending a tissue atlas to additional donors requires assessing annotation performance on donors not represented during classifier training, even when the sequencing platform remains unchanged. When both within-cohort prediction and cross-donor transfer are of interest, reporting both evaluation schemes may be informative, provided they are interpreted as answering different questions.

Several limitations should be considered: we are only considering supervised cell-type classification, a common logistic regression framework, and a finite set of cohorts and latent representations. Donor-held-out cross-validation also captures only one form of dataset shift and does not represent transfer across protocols or independent populations. Expression representations were constructed once for each benchmark cohort rather than re-estimated within individual cross-validation folds. Held-out donors could therefore contribute to representation construction even though their cells were excluded from classifier training. Our comparison isolates the effect of classifier train-test partitioning conditional on these fixed representations, rather than evaluating an end-to-end pipeline trained without access to test donors. Nevertheless, the consistency of the evaluation gap across two distinct datasets with intermediate classification difficulty and multiple representations supports a broader principle for single-cell benchmarking: the train-test split should match the level over which generalization is being claimed.

## 4. Methods

### 4.1 Cohorts, preprocessing, and expression representations

We assembled five publicly available scRNA-seq cohorts spanning peripheral blood, lung, kidney, and pancreas and applied dataset-specific filtering to construct donor-aware cell-type classification benchmarks. The PBMC cohort was obtained from the processed Stephenson et al. dataset distributed through pertpy [5]. The Lung Atlas was derived from the Human Lung Cell Atlas of Sikkema et al. [13], the Kidney cohort from McEvoy et al. [14], the Pancreas cohort from the non-diabetic subset of Bandesh et al. [15], and the Blood Atlas from Terekhova et al. [16].

The PBMC cohort contained 10,107 cells from 23 donors and 13 retained cell types. The Lung Atlas contained 83,966 cells from 104 donors and 31 cell types after restricting to healthy samples and well-supported annotations. Kidney and Pancreas contained 8,746 cells from 19 donors and 8,500 cells from 17 donors, respectively, with nine retained cell types in each. The Blood Atlas benchmark contained 108,682 cells from 166 donors and seven major cell types.

For Lung, Kidney, and Pancreas, count matrices were normalized to 10,000 counts per cell and log-transformed. PBMC retained the processed expression matrix provided with the source data and its existing scVI embedding. Blood Atlas retained the provided transformed expression matrix because gene-level raw counts were unavailable.

We evaluated HVG, PCA, Harmony, and scVI representations where available. We selected 1,000 HVGs, computed 50 principal components, and retained the first 15 for classification. Harmony was applied to the PCA representation using donor identity for PBMC, Kidney, and Pancreas, and source dataset for Lung. scVI embeddings were trained for Lung, Kidney, and Pancreas, while the existing embedding for PBMC was reused. Blood Atlas was evaluated with HVG, PCA, and Harmony only (scVI embeddings cannot be computed as raw counts were unavailable). Representations were constructed once for each fixed benchmark cohort and held constant across evaluation strategies so that random cell-level and donor-held-out comparisons differed only in classifier train-test partitioning.

### 4.2 Training, cross-validation, evaluation design

Cell types were classified using multinomial logistic regression with standardized inputs. Feature scaling was fit on each training partition and applied to the corresponding test partition. The same classifier and hyperparameters were used across datasets, representations, and evaluation strategies.

We compared random cell-level splitting with donor-held-out cross validation. For random evaluation, cells were stratified by cell type into 80% training and 20% test sets, allowing cells from the same donor to appear in both sets. This procedure was repeated across 5 random seeds. For donor-held-out evaluation, donors were partitioned into five folds, with all cells from each test donor excluded from model training. Cell types absent from a training partition were excluded from evaluation for that split.

The primary performance metric was macro F1, which gives each retained cell type equal weight. Accuracy and per-class performance metrics were recorded. Performance was summarized across 5 random splits or donor-held-out folds, and evaluation gap was defined as the difference in mean macro F1 between random cell-level and donor-held-out evaluation.

### 4.3 Sensitivity analyses and settings of contrasting task difficulty

To assess whether measured technical heterogeneity explained differences between evaluation strategies, we repeated classification after adding the collection site (for PBMC) or source dataset (for Lung) as a one-hot-encoded classifier covariate. Donor heterogeneity was also modeled using a binomial generalized linear mixed model of cell-level classification correctness, with evaluation strategy as a fixed effect and donor as a random intercept.

We assessed the effect of training-cohort diversity by repeatedly subsampling different numbers of training donors and evaluating performance on all remaining donors. For PBMC, we additionally performed bidirectional transfer between the Cambridge and Newcastle cohorts. We repeated the analysis after matching cell-type support by downsampling each cell type to the smaller site-specific cell count, and after matching donor counts using equal numbers of donors from the two sites.

Kidney, Pancreas, and Blood Atlas served as highly separable classification settings and were analyzed using the same evaluation framework. For the randomized-label setting, PBMC cell-type labels were replaced with 13 donor-balanced random pseudo-classes while preserving expression profiles and representations. Ten independent label randomizations were evaluated under both splitting strategies, with balanced chance macro F1 equal to 1/13.

## 5. Data Availability, Acknowledgements, Author Contributions

Raw datasets analyzed in this project are all publicly available. Additionally, we have deposited all processed datasets at Zenodo at https://doi.org/10.5281/zenodo.19717558/. All code and data analyses are available on GitHub at https://github.com/danielchen05/scRNA-cross-donor-generalization/.

X.C. participated in project conceptualization, data curation, formal analysis, investigation, methodology, data visualization, and writing. J.L. participated in data curation and writing. A.B. participated in project conceptualization, supervision, funding acquisition, and writing.

This work was supported by the National Institutes of Health/National Institute of General Medical Sciences [R35GM139580 to A.B.]. A.B. is a current employee of insitro, Inc, a shareholder in Alphabet, Inc and is a co-founder and shareholder of CellCipher, Inc. Other authors declare that they have no competing interests.

## Supplementary Figures

**Figure S1:**
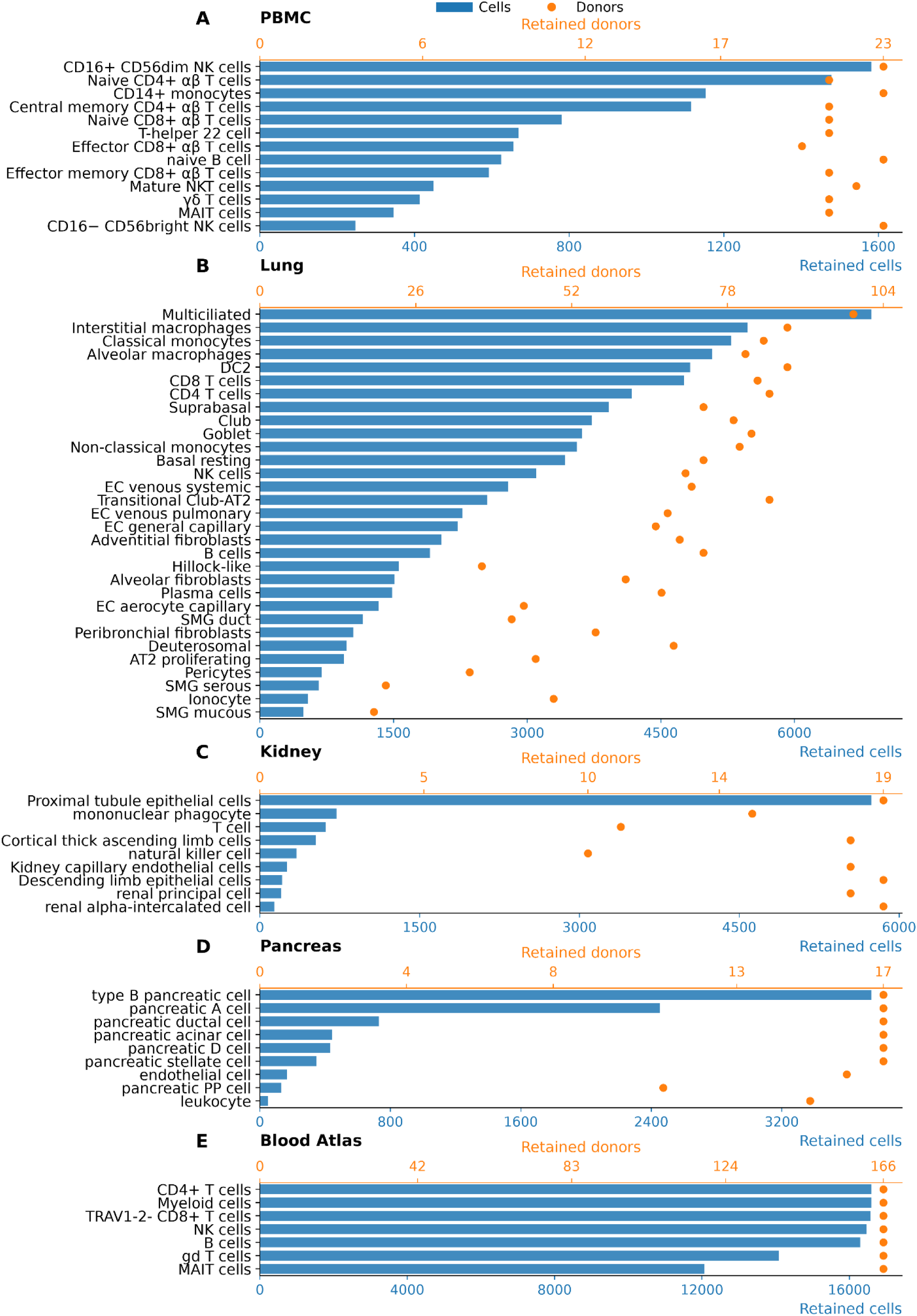
Cell type composition and donor representation across benchmark cohorts after preprocessing. (A): PBMC cohort. Retained cell counts and numbers of contributing donors are shown for each cell type. (B): Lung cohort. Retained cell counts and numbers of contributing donors are shown for each cell type. (C): Kidney cohort. Retained cell counts and numbers of contributing donors are shown for each cell type. (D): Pancreas cohort. Retained cell counts and numbers of contributing donors are shown for each cell type. (E): Blood Atlas cohort. Retained cell counts and numbers of contributing donors are shown for each cell type. Horizontal bars show retained cell counts using the bottom axis, and points show the number of contributing donors using the top axis. Cell types are ordered by decreasing retained cell count within each cohort.

**Figure S2:**
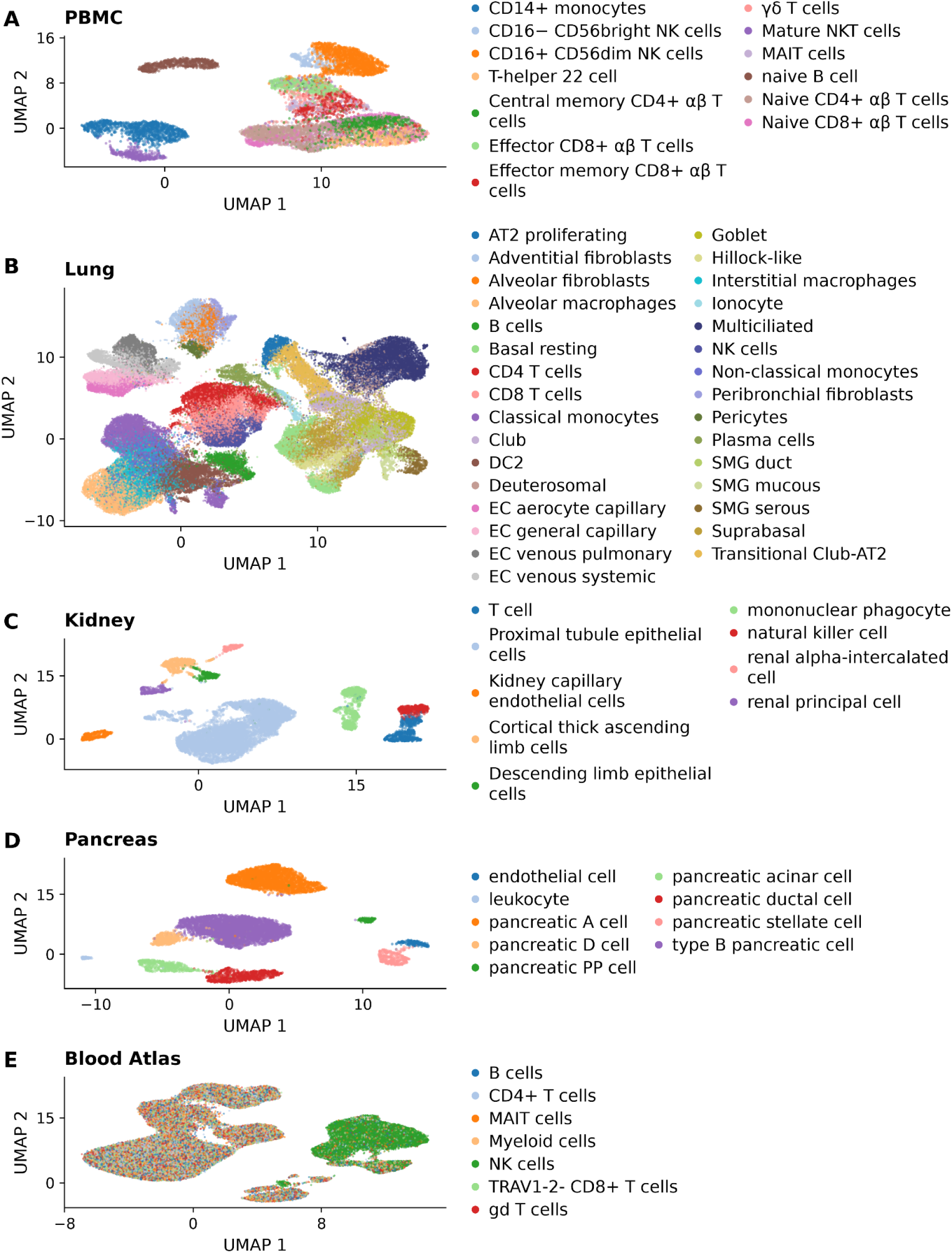
Low-dimensional visualization of retained cell populations across benchmark cohorts. (A): UMAP of the PBMC cohort, colored by annotated cell type. (B): UMAP of the lung cohort, colored by annotated cell type. (C): UMAP of the kidney cohort, colored by annotated cell type. (D): UMAP of the pancreas cohort, colored by annotated cell type. (E): UMAP of the Blood Atlas cohort, colored by annotated cell type. Each point represents one retained cell. UMAP embeddings were constructed from PCA representations independently for each cohort.

**Figure S3:**
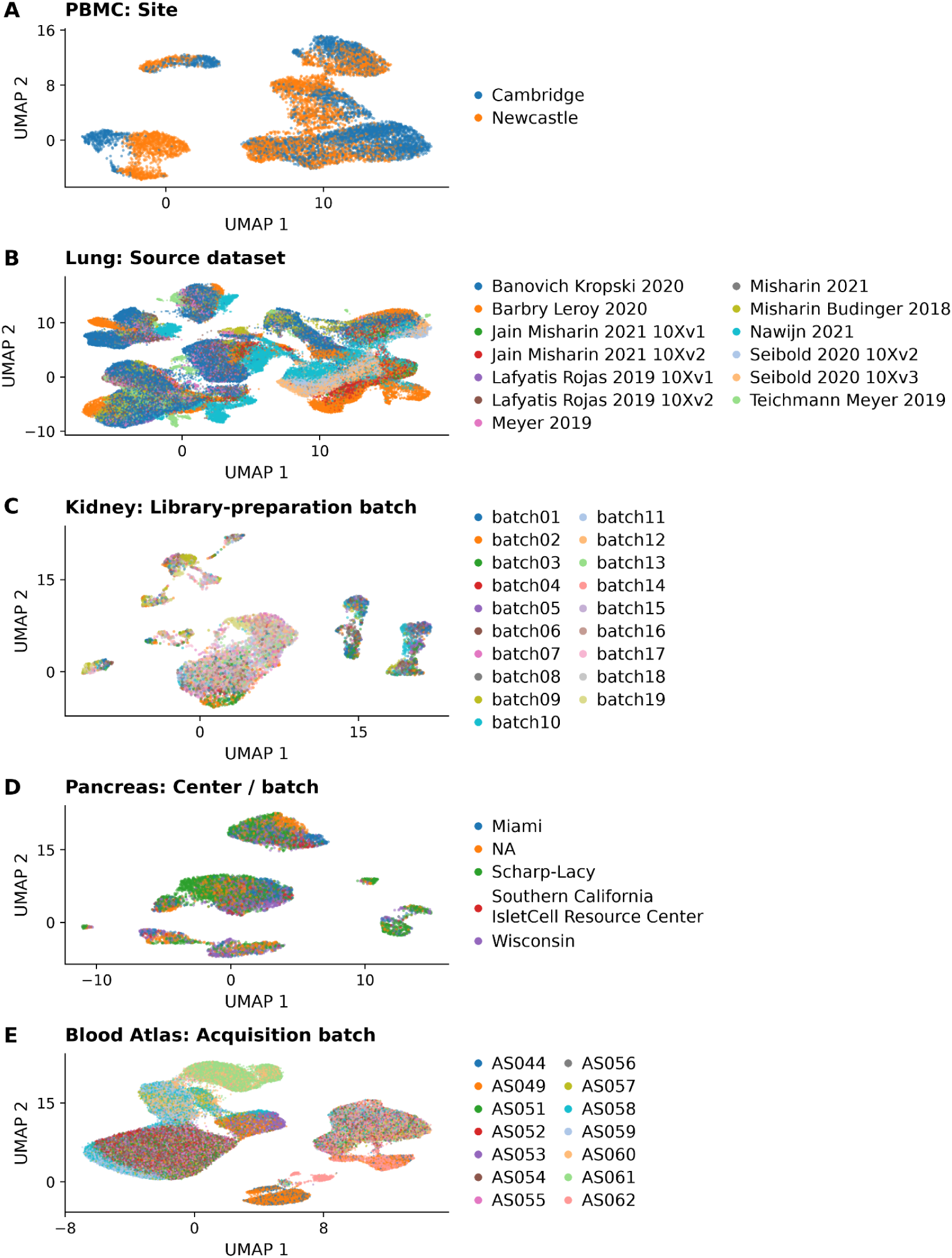
Technical and source-associated structure across benchmark cohorts. (A): PBMC UMAP colored by study site. (B): Lung UMAP colored by source dataset. (C): Kidney UMAP colored by library-preparation batch. (D): Pancreas UMAP colored by contributing center or batch. (E): Blood Atlas UMAP colored by acquisition batch. Each point represents one retained cell, using the same cohort-specific UMAP coordinates as Figure S2.

**Figure S4:**
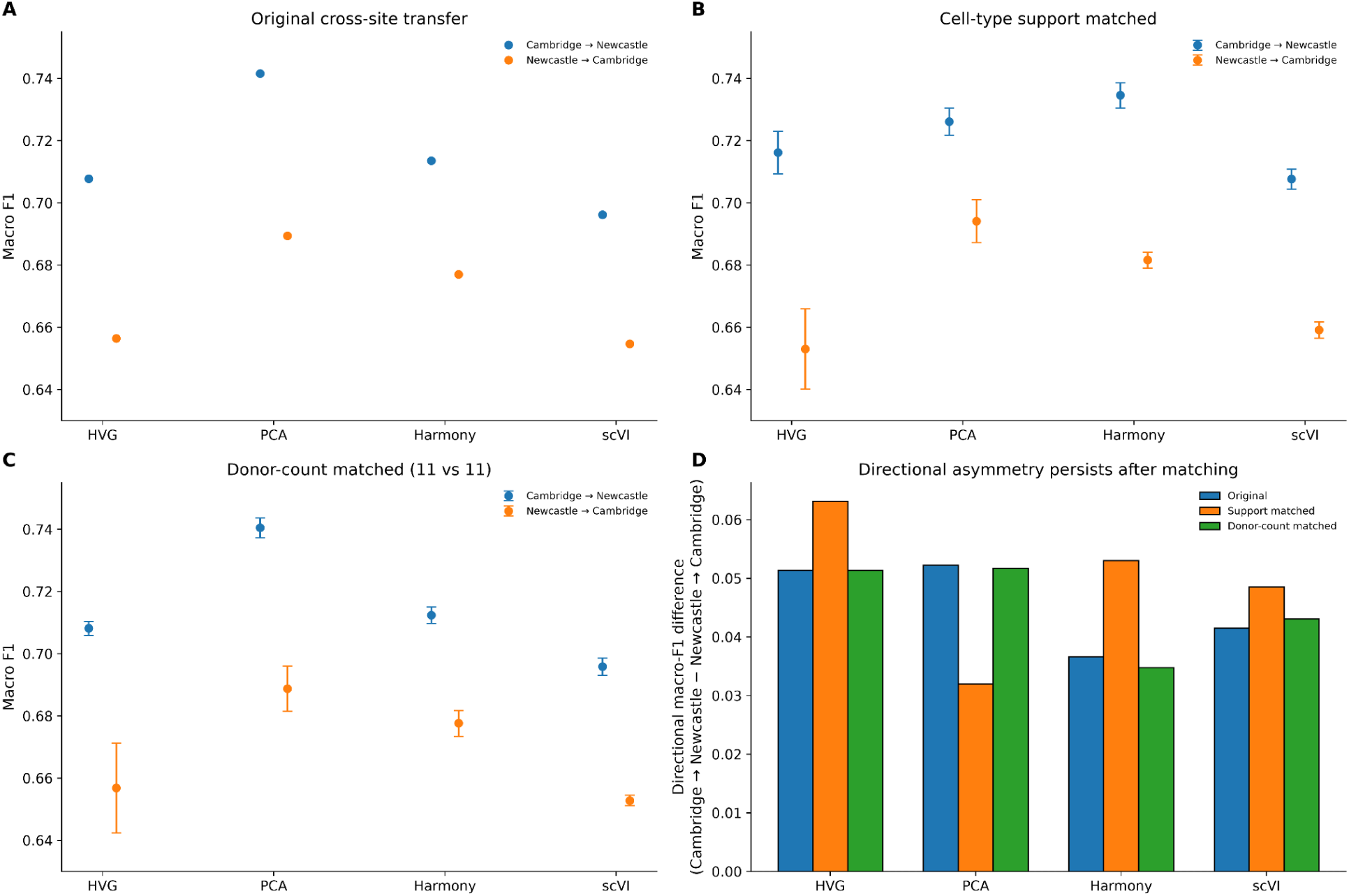
Cross-site transfer in the PBMC cohort shows persistent directional asymmetry after matching analyses. (A): Macro F1 for cross-site transfer between Cambridge and Newcastle across expression representations using the original cohorts. (B): Cross-site transfer after matching cell-type support by downsampling each cell type within each site to the smaller of its two site-specific cell counts. Points and error bars show mean macro F1 ± SD across ten matched resamples. (C): Cross-site transfer after matching donor counts to eleven donors per site; points and error bars show mean macro F1 ± SD across ten donor subsamples. (D): Directional macro F1 difference, calculated as Cambridge-to-Newcastle performance minus Newcastle-to-Cambridge performance, for the original, cell-type-support-matched, and donor-count-matched analyses. Positive values indicate higher performance when training on Cambridge and testing on Newcastle.

**Figure S5:**
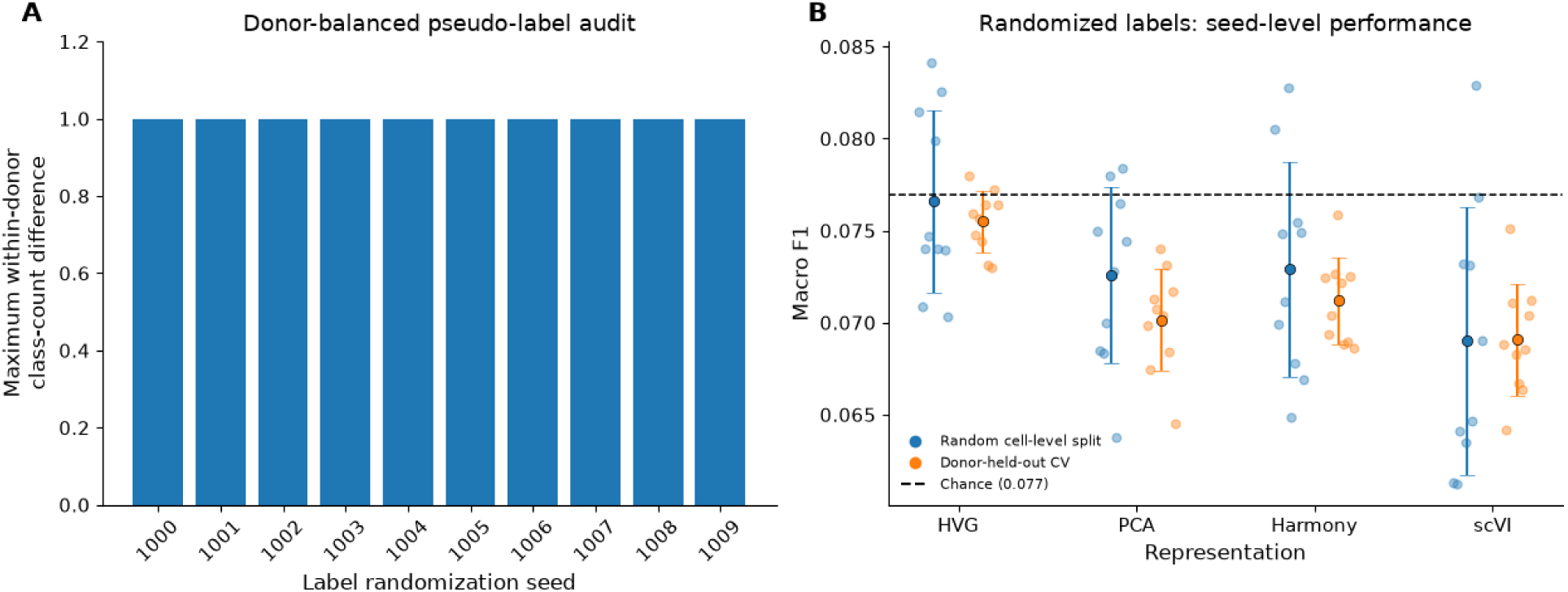
Donor-balanced pseudo-label assignment and seed-level performance in the randomized-label PBMC setting. (A): Maximum within-donor pseudo-class count difference for each of ten label-randomization seeds, calculated as the largest difference between the most and least abundant pseudo-classes within any donor. The maximum is one cell for every seed. (B): Macro F1 for each label randomization under random cell-level splitting and donor-held-out cross-validation across expression representations. Each light point represents one label randomization, evaluated using one random cell-level split or the mean across five donor-held-out folds. Darker points and error bars show the mean ± SD across the ten label randomizations. The dashed line marks the balanced-chance reference of 1/13, approximately 0.077.

## Notes

https://github.com/danielchen05/scRNA-cross-donor-generalization

https://doi.org/10.5281/zenodo.19717558

### Competing Interest Statement

A.B. was supported by NIH/NIGMS R35GM139580. A.B. is a current employee of insitro, Inc, a shareholder in Alphabet, Inc and is a co-founder and shareholder of CellCipher, Inc. Other authors declare that they have no competing interests.

### Summary of Updates

Revised the manuscript to emphasize classification task difficulty and separability, replacing ceiling/floor terminology. Clarified the study motivation and practical implications. Updated figures and captions, and revised data availability information. The main results and conclusions remain unchanged.

